# Theoretical estimate of the effective pKa of titratable lipids using continuum electrostatics

**DOI:** 10.64898/2026.04.06.716676

**Authors:** Sreyoshi Sur, Alan Grossfield

## Abstract

The apparent pKa of ionizable lipids in lipid nanoparticles (LNPs) is a key determinant of RNA encapsulation during formulation and endosomal release after cellular uptake. However, it is difficult to predict the effective pKa of a given ionizable lipid solely from its solution pKa, because it is sensitive to the membrane’s composition, as well as solution conditions such as the salt concentration. We developed a simple continuum electrostatics model, based on Gouy-Chapman theory, to predict the shift in effective pKa for ionizable lipids in lipid bilayers as a function of salt concentration and membrane composition. We derive equations for the surface potential and fraction of lipids charged, which are solved self-consistently as a function of solution pH to extract the titration curve and effective pKa. The model shows that the shift in effective pKa is largest when the concentration of titratable lipid is high, and the effect is diminished by increasing salt concentration. We provide a python implementation of the model and an interactive notebook that will allow users to further easily explore the predicted pKa shifts as a function of formulation variables.

## 1 Introduction

Lipid nanoparticles (LNPs) are used to deliver custom siRNA and mRNA molecules for gene silencing and protein translation, respectively, among other applications. ^1,2^. Although this technology was developed over decades, it became part of the popular imagination during the COVID19 pandemic, when synthetic lipid nanoparticles were used to deliver the mRNA vaccines developed by Moderna and Pfizer. ^2^ While best known for its use in vaccines, nanoparticle-based RNA delivery can also be used for protein replacement, gene editing, and cancer immunotherapy. ^2,3^

Nanoparticles that effectively encapsulate and deliver RNA generally contain a precise, carefully calibrated mixture of lipids. This includes titratable cationic lipids, which are crucial for RNA encapsulation under acidic conditions, and phospholipids, which help the nanoparticles maintain structural integrity. Cholesterol also stabilizes nanoparticles structure, while polyethylene glycol (PEG) lipids are often added to increase circulation time in the bloodstream and improve delivery ^3^.

The ionizable lipids used in synthetic lipid nanoparticles are unusual in that they are cationic at low pH and neutral at physiological pHs or higher, where natural lipids are more commonly zwitterionic or anionic. As a result, under acidic conditions during formulation, these lipids help stabilize binding and encapsulation of negatively charged nucleic acids, while at physiological pH the neutral lipids form an interior phase that maintains a stable nanoparticle structure. Upon successful delivery to target cells, the acidic environment of the endosome recharges the titratable lipids, disrupting the nanoparticle structure and releasing the RNA. ^2,4,5^

Formulating nanoparticles that efficiently bind and deliver RNA is highly sensitive to both the composition of the lipid mixture and the choice of pH during formulation, and identifying optimal conditions is a labor-intensive process that involves a lot of trial and error. ^6,7^ If we are to rationally design better lipid nanoparticles, we need to understand the physical factors governing their structure and stability, and in particular we must understand the factors controlling the charge state of the titratable lipids.

The obvious choice to begin this process is to measure the pKa of the relevant chemical moieties in solution, e.g. the tertiary amine found in lipids such as KC2. ^8,9^ However, while a solution measurement is straightforward — the pKa of the tertiary amine is around 9-10 in aqueous solution ^10^ — the pKa changes by as much as 2-4 units when incorporated into an LNP. Several physical factors are involved in this shift, including changes in hydration, interactions with the surrounding phospholipids, and the electrostatic repulsion of protons due to other ionizable lipids in the nanoparticle, so the shift in pKa is sensitive to the composition of the LNP ^7,10–13^.

Recently, several groups have attempted to measure the shifted pKa using molecular dynamics simulations, including umbrella sampling ^14^ and constant-pH molecular dynamics ^15,16^. However, these calculations each treat a single set of conditions and are computationally expensive, so it would be desirable to have a simpler theoretical framework to understand pKa shifts.

Here, we present a theoretical treatment of the apparent pKa of ionizable lipids in LNPs, based on continuum electrostatics. Combining Gouy-Chapman theory (essentially, the Poisson-Boltzmann equation for a charged surface) ^17–20^ with an equilibrium treatment of the protonation state of a titratable lipid, we self-consistently solve for the fraction of charged lipids as a function of solution pH, and thus extract the apparent pKa as a function of lipid composition and salt concentration.

## 2 Methods

### 2.1 Derivation

The basic strategy used in developing this model is to represent the equilibrium between charged and neutral forms of the titratable lipid using continuum electrostatics; the presence of the cationic form produces a net charge at the membrane interface, which in turn shifts the effective pKa. We begin the derivation by recapitulating the derivation of Gouy-Chapman theory. ^18–21^ The Poisson-Boltzmann equation in one dimension is:

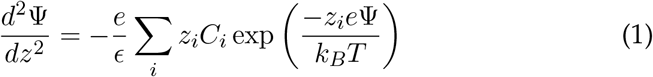

where Ψ is the electrostatic potential, *z* is the distance from the membrane interface, *e* is the charge on an electron, *ϵ* is the dielectric constant, *z*_*i*_ is the valence of ion *i, C*_*i*_ is the concentration of ion *i* in solution, *k*_*B*_ is Boltzmann’s constant, and *T* is the temperature.

If we restrict ourselves to the case of a 1:1 electrolyte, the sum simplifies to:

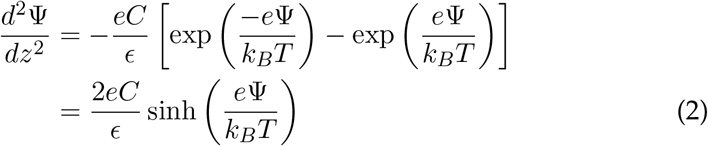

where *C* is the concentration of electrolyte.

One can solve this differential equation analytically with the boundary conditions that the potential and electric field go to zero as *z* → ∞far from the surface, which yields

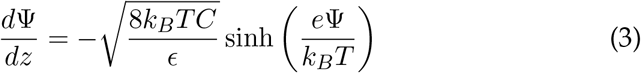

We can then apply Gauss’ law to get an expression for the electrostatic potential

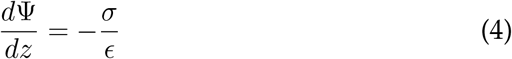

Combining Equations 3 and 4 gives:

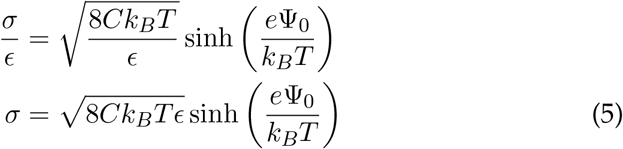

In most cases where Gouy-Chapman theory is applied, the charge density *σ* is a constant property of the surface. However, for a lipid bilayer containing titratable lipids, the charge depends on the pH of the solution. Specifically, the average charge density is the product of the area density of the titratable lipids and the fraction of these lipids that are charged. With this in mind, we rearrange Equation 5 to express the surface potential in terms of the charge density:

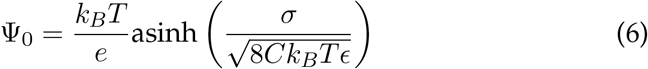

where Ψ_0_ is the surface potential at *z* = 0.

To estimate the charge density on the surface, we next consider the equilibrium between the charged and neutral forms for the ionizable lipid. For purposes of this discussion, we will assume the lipid is either cationic or neutral and that solvation in the membrane-bound form is similar to that in solution. In this case, the free energy change upon charging can be written as:

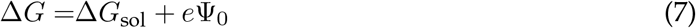

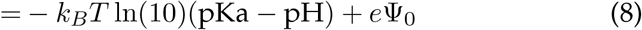

where Δ*G*_sol_ is the free energy change upon charging in solution, and the second term is the work required to bring a charge from infinity to the surface against the electrostatic potential. In this case, the probability that a lipid is in the charged state is

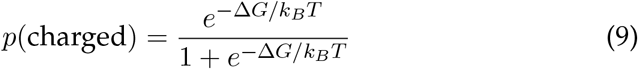

We can approximate the area density as the weighted average of the areas per lipid for all species in the membrane:

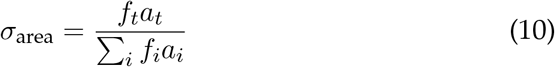

where *a*_*i*_ is the area per lipid of species *i, f*_*i*_ is the mole fraction of that species in the bilayer, and the subscript *t* refers to the titratable species. Thus, the total charge density is:

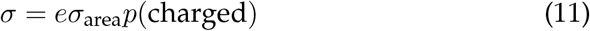

### 2.2 Implementation

Equations 6 and 11 can be solved numerically using an iterative approach. We begin with an initial guess for the surface potential, calculate the charge density using Equation 11, then substitute it into Equation 6 to get a new surface potential, and continue iterating until convergence, which we define as an iteration where the fractional change in surface charge density is less the 10^−8^. To improve numerical stability, a predictor-corrector scheme was used; otherwise, iteration may fail to converge for some combinations of parameters. This approach was used to compute the fraction of charged lipids as a function of pH for a given set of conditions. To compute the effective pKa, we apply cubic spline interpolation to identify the pH where half of the titratable lipids are charged.

A python implementation of the method is available at https://github.com/GrossfieldLab/pka-theory-paper-public. An interactive version is available at https://colab.research.google.com/drive/1o0VDov8yNagc9MjXrTLB9MWgwL6xY8UM

## 3 Results

Solving equations 6 and 11 self-consistently, we can compute the fraction of titratable lipids that are charged as a function of the buffer pH, from which we can read off the effective pKa.

Figure 1 shows the effect of varying the salt concentration on the effective pKa of the titratable lipids. The curves shift to the right as the salt concentration increases, indicating an increase in the effective pKa; in the limit of very high salt concentration, the titration curve approaches that expected in solution. Physically, this can be understood as the result of electrostatic screening by the salt reducing the electrostatic potential at the surface, thus making it easier to protonate the lipids and increasing the effective pKa. At the lowest salt concentration considered (0.1 M), the effective pKa is shifted by roughly 1 pH unit.

**Figure 1:**
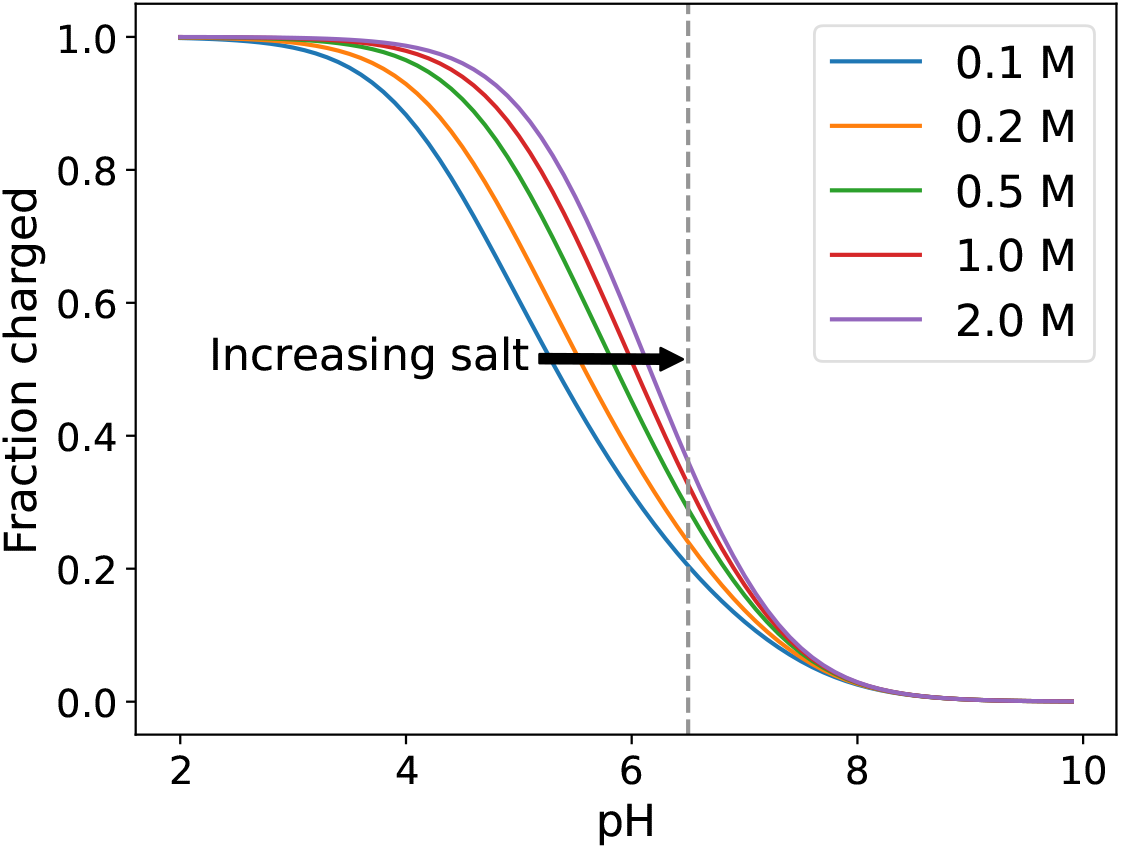
Dependence of the fraction of lipids charged on the salt concentration. The vertical dashed line indicates the solution pKa of the titratable lipid is 6.5, the area per lipid is 60 Å^2^, the temperature is 300 K, and the mole fraction of the titratable lipid is 0.5. The key indicates the salt concentration in molar units.

Along the same lines, Figure 2 shows the model’s predictions regarding the effects of the bilayer’s composition, specifically the mole fraction of titratable lipids. As the fraction of titratable lipids in the membrane increases, the curves shift to the left, indicating a decrease in the effective pKa. We interpret this as the result of increased charge density at the molecular surface, which in turn increases the electrostatic potential and thus the work required to bring a proton to the surface.

**Figure 2:**
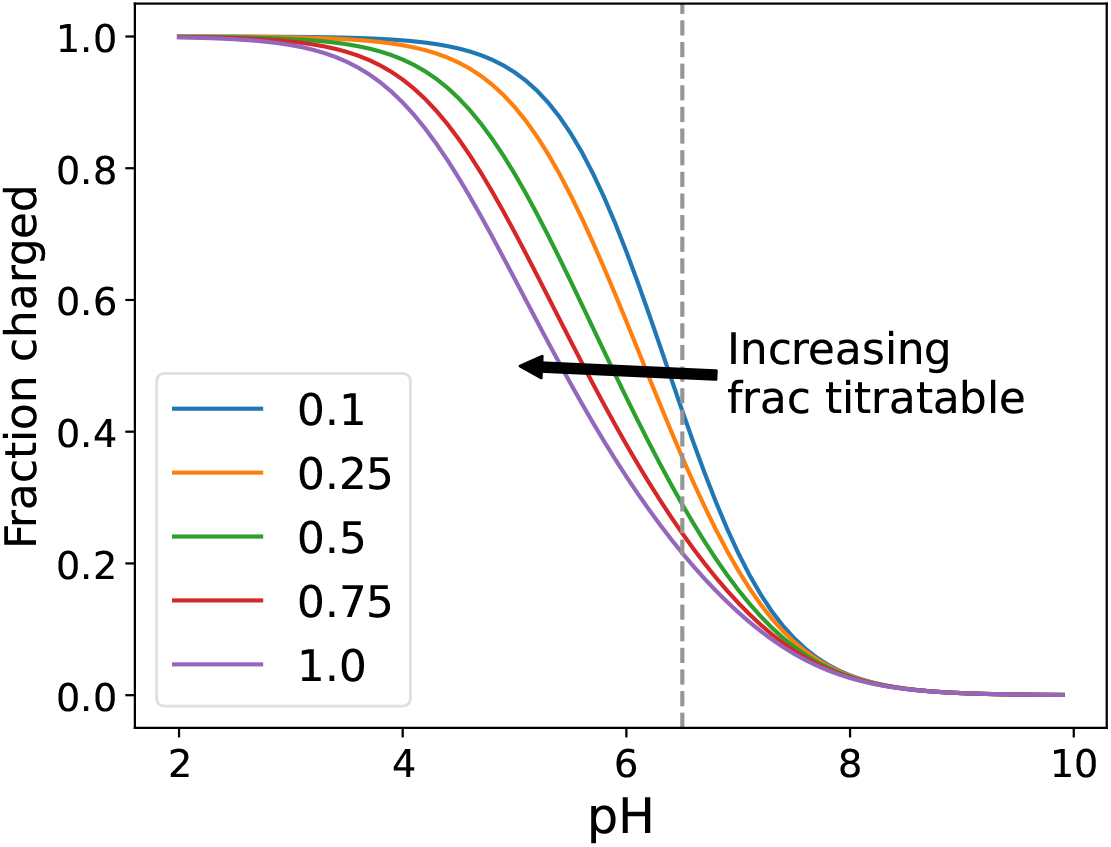
Dependence of the fraction of lipids charged on the concentration of titratable lipids. The intrinsic pKa is 6.5, the area per lipid is 60 Å^2^, the temperature is 300 K, and the salt concentration is 500 mM. The key indicates the mole fraction of titratable lipids in the bilayer.

Figure 3 shows the combined effects of salt and membrane composition on lipid titration by mapping the effective pKa. As expected, the pKa decreases as the mole fraction of titratable lipids increases or the salt concentration decreases. The largest amplitude pKa shift, roughly -1.7 pH units, occurs when the membrane is entirely composed of titratable lipids and the salt concentration is 0.1 M.

**Figure 3:**
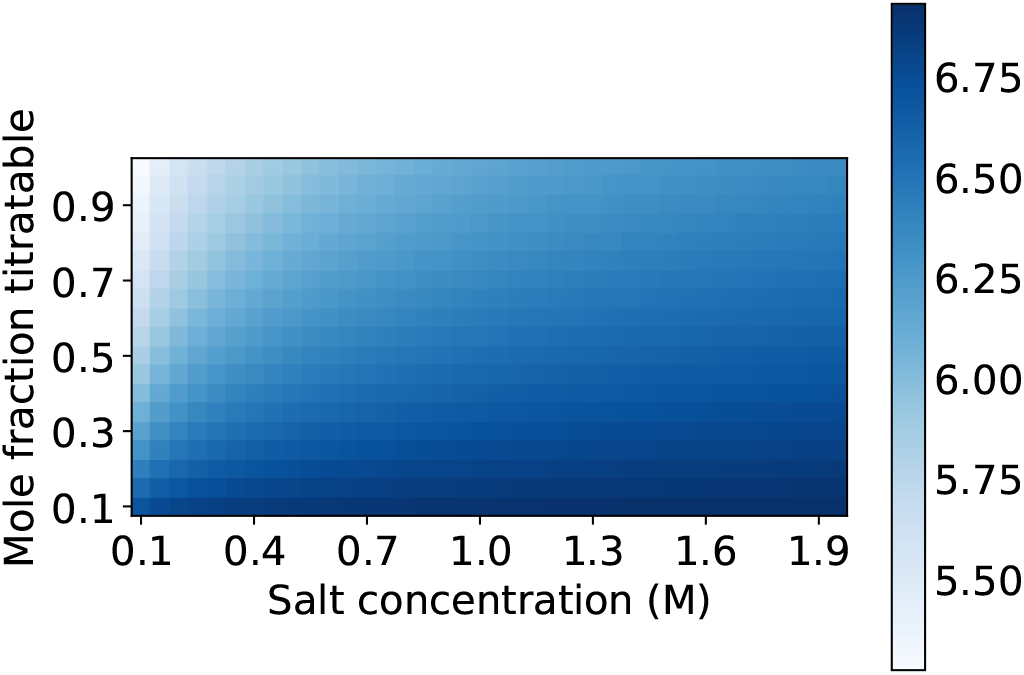
Effective pKa as a function of the mole fraction of titratable lipid and salt concentration. The intrinsic pKa is 6.5, the temperature is 300 K, and the area per lipid is 60 Å^2^.

## 4 Discussion

The proposed model makes a number of assumptions, which necessarily limit its quantitative accuracy and ability to predict the properties of any specific system; the value of the model is in its ability to yield intuition about the trends expected for a typical system. That said, it is worth explicitly laying out these assumptions, discussing their implications, and explaining how they might be relaxed.

The largest assumption is the applicability of continuum electrostatics, and the Gouy-Chapman theory in particular, to the electrostatics of a lipid bilayer. In particular, this model assumes that charge is distributed uniformly on the surface, solvation can be treated using a simple dielectric, and salt obeys the Poisson-Boltzmann equation. These assumptions are essential to the entire approach, and cannot easily be relaxed. In particular, the problem can only be reduced to 1 dimension if the charge density is uniform.

The assumption that the salt is a 1:1 electrolyte (Equation 2) is similarly essential, since it transforms the solution from a sum over ionic species to a pair of terms identical except for the sign of the charge that can be reduced to a single sinh term. Without this simplification, the inversion to get the surface potential (Equation 6) is not possible.

The derivation explicitly treats the bilayer as a plane, which is only appropriate when considering a large nanoparticle (or rather, a large vesicle, which is the usual form for these lipids under acidic conditions). The Poisson-Boltzmann equation can be solved analytically for other geometries, such as a cylinder or sphere, but it is not clear whether the approach described in this paper would still be tractable. We do not regard this assumption as a major limitation, since it is hard to see how bilayer curvature would qualitatively change the results except for very small vesicles.

We additionally assume that in the limit of very low concentrations of titratable lipid, the effective pKa will be essentially the same as in solution; this is most easily seen by considering the lower edge of the effective pKa plot in Figure 3. This is implicit in the simplistic continuum treatment of the membrane, and neglects electrostatic effects from the bilayer dipole potential from the non-titratable lipids, as well as any potential differences in the solvation of the titratable lipid’s headgroup when in the bilayer vs in solution. The dipole potential could in principle be included as an additional term in the electrostatic potential, either as an additional constant in Equation 6 or more correctly as part of Equation 3. However, doing so would introduce an additional free parameter to the model but appears unlikely to change the qualitative behavior. Similarly, changes to headgroup solvation could be easily treated with an additional free energy term in Equation 8 or equivalently as a constant shift in the solution pKa. This does not seem worthwhile, given the qualitative goals of the model.

Although the model as currently presented only handles a single titratable species, it should be straightforward to extend it to multiple titratable species. In this case, Equation 8 would define a pKa and Δ*G* for each species, and thus each species would have its own probability of being charged, calculated with Equation 9. Computing the charge density would require replacing Equation 11 with a sum over species.

For the purposes of the model, we define the maximum charge density for the membrane surface as the area density of the titratable lipid multiplied by 1 electron charge. The area density of the titratable lipid is estimated using Equation 10, which assumes that areas per lipid are additive. Cholesterol, a common component of lipid nanoparticles, has a condensing effect of liquid crystalline phospholipids, which clearly violates the additivity assumption ^22,23^. However, the deviations from additivity are generally only a few percent, so this is unlikely to be a major source of error in the model.

In the same vein, we assume that the lipid areas are the same for the charged and neutral forms of the titratable lipid. Short of performing careful experimental measurements or atomistic simulations, this assumption is difficult to evaluate. However, a more significant challenge is that the neutral form of the titratable lipids is able to partition into the interior of the bilayer; indeed, this is believed to be the mechanism by which the vesicles formed at low pH transform into monolayer-coated nanoparticles at neutral or higher pHs.

Integrating partitioning into our model will be far more challenging, since we would have to allow for a global change in morphology. As lipids partition into the interior, the area of the leaflet will decrease. At some point, the area of the leaflet will decrease to the point where it can no longer enclose the interior, and the system will transition from a bilayer to a monolayer-coated nanoparticle. Computing the free energy for the system in a continuum model would require accounting for the intrinsic preference of the neutral form to partition into the interior, the density of the interior, and the area per lipid of the remaining lipids and cholesterol, as well as the area modulus of the leaflet and the compressibility of the interior. While we suspect this is achievable, it is beyond the scope of the present work.

## 5 Conclusion

We described a simple continuum electrostatics model to describe the effects of lipid composition and salt concentration on the effective pKa of titratable lipids in lipid bilayers and nanoparticles. The effective pKa is crucial in determining the properties of LNPs and their ability to effectively encapsulate and deliver RNA, and moving beyond empirical tuning of the formulation process would be a significant step forward towards rational design of better LNPs. Although it lacks details of specific lipid chemistry, this theory provides a simple framework for thinking about how formulation variables might affect the properties of the resulting LNP.

## 6 Acknowledgements

We thank Akshara Sharma for helpful discussions and comments on the manuscript.

